# From genetic disposition to academic achievement: The mediating role of non-cognitive skills across development

**DOI:** 10.1101/2025.02.27.640510

**Authors:** Quan Zhou, Wangjingyi Liao, Andrea G. Allegrini, Kaili Rimfeld, Jasmin Wertz, Tim Morris, Laurel Raffington, Robert Plomin, Margherita Malanchini

## Abstract

Genetic effects on academic achievement are likely to capture environmental, developmental, and psychological processes. How these processes contribute to translating genetic dispositions into observed academic achievement remains critically under-investigated. Here, we examined the role of non-cognitive skills—e.g., motivation, attitudes and self-regulation—in mediating education-associated genetic effects on academic achievement across development. Data were collected from 5,016 children enrolled in the Twins Early Development Study at ages 7, 9, 12, and 16, as well as their parents and teachers. We found that non-cognitive skills mediated polygenic score effects on academic achievement across development, and longitudinally, accounting for up to 64% of the total effects. Within-family analyses highlighted the contribution of non-cognitive skills beyond genetic, environmental and demographic factors that are shared between siblings, accounting for up to 83% of the total mediation effect, likely reflecting evocative/active gene-environment correlation. Our results underscore the role of non-cognitive skills in academic development in how children evoke and select experiences that align with their genetic propensity.

## Introduction

Academic achievement, commonly measured as school grades during childhood and adolescence, is associated with a range of positive life outcomes, including better physical and mental health^1,2^ and higher income^3–5^. Research indicates that genetic differences between people partly explain variations in academic achievement, a concept known as heritability^6^. Twin studies, which estimate the relative contribution of genetic and environmental factors by comparing identical and fraternal twins, have shown that 40 to 60% of the differences in educational attainment and academic achievement can be attributed to genetic factors^7,8^.

DNA-based methods have also quantified academic achievement, estimating heritability at 20 to 30% during childhood and adolescence^7,9,10^. Another way of capturing education-associated genetic effects is to construct a polygenic score (PGS)^11^. Polygenic scores aggregate the effects of numerous genetic variants identified in large-scale genome-wide association studies, creating a composite index that captures known genetic effects on a trait^12^. Education-associated polygenic scores have been found to account for 12 to 16% of the variance in adult educational attainment (years of schooling)^13^. In addition, a polygenic score combining education^14^ and cognition-associated genetic variants was found to account for 15% of the variance in academic achievement at the end of compulsory education in a UK-based sample^15^, a figure comparable to other well-established predictors of academic achievement, such as family socioeconomic status^16^.

Genetic effects on academic achievement likely encompass a combination of environmental, developmental, and psychological processes. To explore this further, two recent studies investigated how family environments mediate these genetic influences. The first study showed that, beyond socioeconomic status, multiple aspects of the family environment – such as supportive parenting, household chaos, and TV consumption – mediated polygenic score associations with academic achievement throughout compulsory education^17^. By employing a sibling-difference design to distinguish genetic associations shared between siblings (between-family effects) and those unique to each child (within-family effects), the study found that family environments were predominantly correlated with between-family genetic effects^17^. These effects likely reflect passive gene-environment correlation, suggesting that children, by growing up with their biological parents, are exposed to environmental experiences that correlate with their genetics^18^. The second study found that mothers’ educational attainment polygenic scores were associated with their children’s educational outcomes, beyond the children’s own genetics. This ‘genetic nurture’^19^ pathway from mother’s genetics to her children’s attainment is thought to capture environmental processes that covary with parental genetic propensities^20^ and was mediated by cognitively stimulating parenting^21^. However, environmental, developmental and psychological experiences other than family environments that may translate genetic propensity into academic achievement remain under-investigated.

The current study investigates the role of non-cognitive skills in mediating education-associated genetic effects on academic achievement across development. The term ‘*non-cognitive skills*’ describes a set of attitudes and characteristics that impact life outcomes beyond what cognitive tests can measure and predict^22^. These skills encompass motivation, perseverance, mindset, learning strategies, and social skills, and self-regulatory strategies^23^. Non-cognitive skills independently contribute to educational success beyond cognitive ability. A study of 16-year-olds found self-efficacy and personality traits significantly influenced academic achievement beyond cognitive ability^24^. Other studies also link personality, self-regulation, and motivation to academic success^25–28^. More recently, our research highlighted that the association between non-cognitive skills and academic achievement substantially increased from ages 7 to 16^29^. Furthermore, previous research showed that greater self-control and, to a lesser extent, interpersonal skills, partly explain the genetic link to educational attainment^30^.

We explored the role of non-cognitive skills in mediating education-genetic effects on academic achievement through five key questions: First, do non-cognitive skills mediate polygenic score associations with academic achievement across compulsory education? Second, can mediation effects be captured even when separating education-associated genetics into a cognitive and a non-cognitive component^29,31^ (i.e., are education-associated cognitive and non-cognitive genetic effects associated with academic achievement via partly distinct non-cognitive mechanisms?)? Third, and importantly, do non-cognitive skills mediate polygenic score-academic achievement associations longitudinally? Fourth, do non-cognitive skills mediate polygenic score associations with academic achievement even when limited to differences within pairs of siblings? And lastly, is the mediating role of non-cognitive skills robust when controlling for other factors known to be related to both academic achievement and non-cognitive skills, namely general cognitive ability and family socio-economic status?

To address these questions, we employed a design with unique features, including the measurement of several non-cognitive skills across development, such as academic motivation, academic interest, and self-regulation at age 7, 9, 12 and 16 (see Methods). Data on non-cognitive skills were obtained from multiple raters, including teacher, parents, and the children themselves. Furthermore, we used a sibling difference design allowing to partly adjust for shared sibling processes, such as passive gene-environment correlation^17,32^, demographic between-family factors, population stratification^33^, and assortative mating^34^. These preregistered analyses (https://osf.io/vmpf7/) provide a detailed exploration of how non-cognitive skills contribute to translating genetic disposition into observed differences between children in academic achievement across compulsory education.

## Results

### Creating latent dimensions of non-cognitive skills

Several measures of non-cognitive skills were collected at different ages, with ratings provided by parents, teachers, and the twins themselves. Following our previous work, two latent dimensions of non-cognitive skills were created using confirmatory factor analyses^29^: education-specific non-cognitive skills (including attitudes towards school, self-perceived academic ability, academic interest and curiosity) and self-regulation (encompassing behavioural and emotional regulation). See Methods for a detailed description.

### Correlations between polygenic scores, non-cognitive skills and academic achievement

Descriptive statistics are presented in **Supplementary Note 1** and **Supplementary Table 1**. All variables were normally distributed; therefore, no transformations were applied prior to analyses.

We first analysed correlations of polygenic scores constructed from prior genome-wide association studies of educational attainment (*EA*^14^), cognitive (*Cog*) and non-cognitive (*NonCog*) skills, operationalised as genetic variants associated with education after accounting for genetic effects associated with cognitive performance^29^, with academic achievement and the latent non-cognitive dimensions rated by parents, teachers and self across compulsory education. These analyses were performed on unrelated individuals by selecting one twin from each pair randomly to account for relatedness. The correlations between polygenic scores and academic achievement at all ages were statistically significant, moderately sized (**Supplementary Figure 1**) and increased across development, particularly for the EA polygenic score (r ranging from 0.25 at age 7 to 0.36 at age 16) and the NonCog polygenic score (r ranging from 0.10 at age 7 to 0.22 at age 16; **Supplementary Figure 1**). The correlations between polygenic scores and latent dimensions of non-cognitive skills were modest and statistically significant, except for self-rated non-cognitive skills at age 9 (**see Supplementary** Figure 2**; Supplementary Table 2**). The correlations between latent dimensions of non-cognitive skills and academic achievements were positive, moderately sized and statistically significant across all ages (r ranging between 0.13 and 0.63) (**Supplementary Figure 2; Supplementary Table 2**). Moderate and positive correlations were also observed when considering individual measures of non-cognitive skills, including academic interest, self-perceived ability, grit, and curiosity (**see Supplementary Table 3**).

### Non-cognitive skills mediate polygenic score effects on academic achievement

We applied mediation models to examine the extent to which latent dimension of non-cognitive skills (i.e., education-specific non-cognitive characteristics and self-regulation) mediated the association between genetic disposition towards education, measured using the educational attainment polygenic score, and academic achievement across compulsory education. **Figure 1** (top panel) depicts the results of the mediation models for the educational attainment polygenic score. Indirect (mediated) effects for self-regulation were developmentally stable, similar across different raters, and small in magnitude, ranging from ß = 0.01 to ß = 0.03. In contrast, indirect (mediated) effects for education-specific non-cognitive skills increased across compulsory education, varied across raters, and were larger in magnitude. When considering self-reported measures, indirect (mediated) effects increased from zero at age 9, to ß = 0.02 [95% CI, 0.01, 0.04] at age 12 to ß = 0.08 [95% CI, 0.06, 0.11] at age 16. The indirect (mediated) effects of education-specific non-cognitive skills at age 9 were larger for parent-rated (ß = 0.05 [95% CI, 0.02, 0.07]) and teacher-rated measures (ß = 0.13 [95% CI, 0.10, **0.15], (**see Fig 1 top panel; Supplementary Table 4).

**Fig 1:**
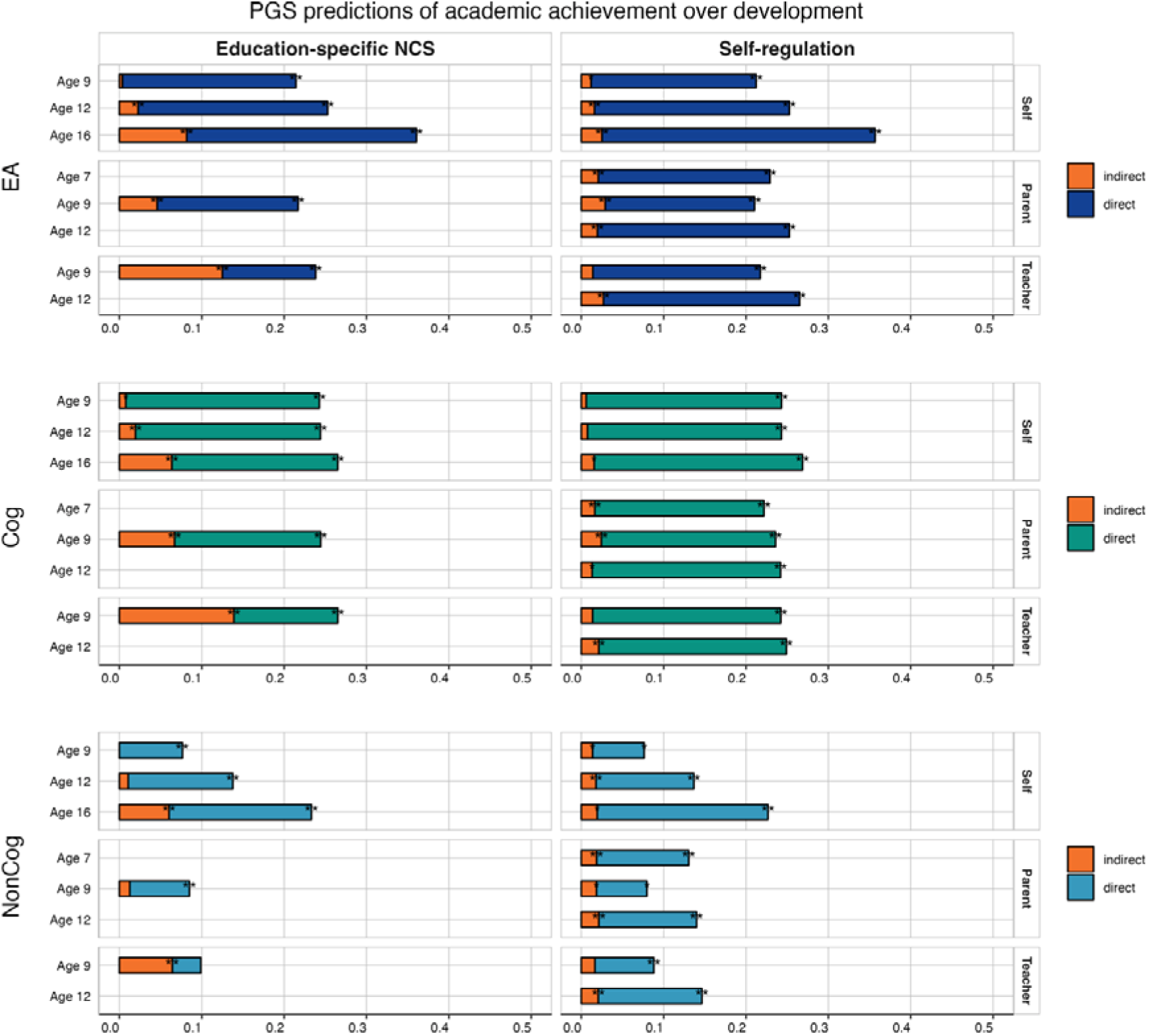
Polygenic score associations with academic achievement mediated by latent dimensions of education-specific non-cognitive skills and domain-general self-regulation. The length of each bar represents the total effect (standardised beta coefficient) between the polygenic scores for educational attainment^14^ (EA; top panel), cognitive^29^ (Cog, middle panel) and non-cognitive^29^ (NonCog; bottom panel) and academic achievement at ages 7, 9, 12, and 16. The orange portion of each bar shows the indirect effect (i.e., mediated by latent factors of non-cognitive skills), while the remaining portion of each bar shows the direct effect of polygenic scores on academic achievement (i.e., not mediated by each non-cognitive measure). The left panels show mediation effects for education-specific non-cognitive skills (NCS), while the right panels show mediation effects associated with self-regulation. * = p < 0.05, ** = p < 0.01 after applying FDR correction. See model estimates in Supplementary **Tables 4-6**.

The two dimensions of education-specific non-cognitive skills and self-regulation also mediated the cognitive and non-cognitive polygenic score-academic achievement association across compulsory education. The strongest mediation effects were observed for teacher-rated education-specific non-cognitive skills at age 9, which accounted for up to 48% of the total cognitive polygenic score prediction, and up to 64% of the total non-cognitive polygenic score prediction (**Fig 1 middle and bottom panels, and Supplementary Tables 5 and 6 for** results on cognitive and non-cognitive polygenic scores obtained using a different model^31^).

### A detailed analysis of education-specific non-cognitive measures as mediators of polygenic score effects on academic achievement

We conducted a more detailed set of analyses to investigate the role played by specific non-cognitive measures in the association between polygenic scores and academic achievements (**Fig 2 and Supplementary Tables 7-9**). Mediation effects were stronger for student-centred non-cognitive variables such as academic interest, academic perceived ability, self-concept and curiosity, as compared to measures capturing broader aspects of the learning environment, such as classroom satisfaction and peer support (**see Fig 2**). In line with the results we obtained when considering latent non-cognitive dimensions, the mediating role of parent-rated and teacher-rated non-cognitive measures at age 9 was stronger than what we observed for self-rated non-cognitive measures.

**Fig 2:**
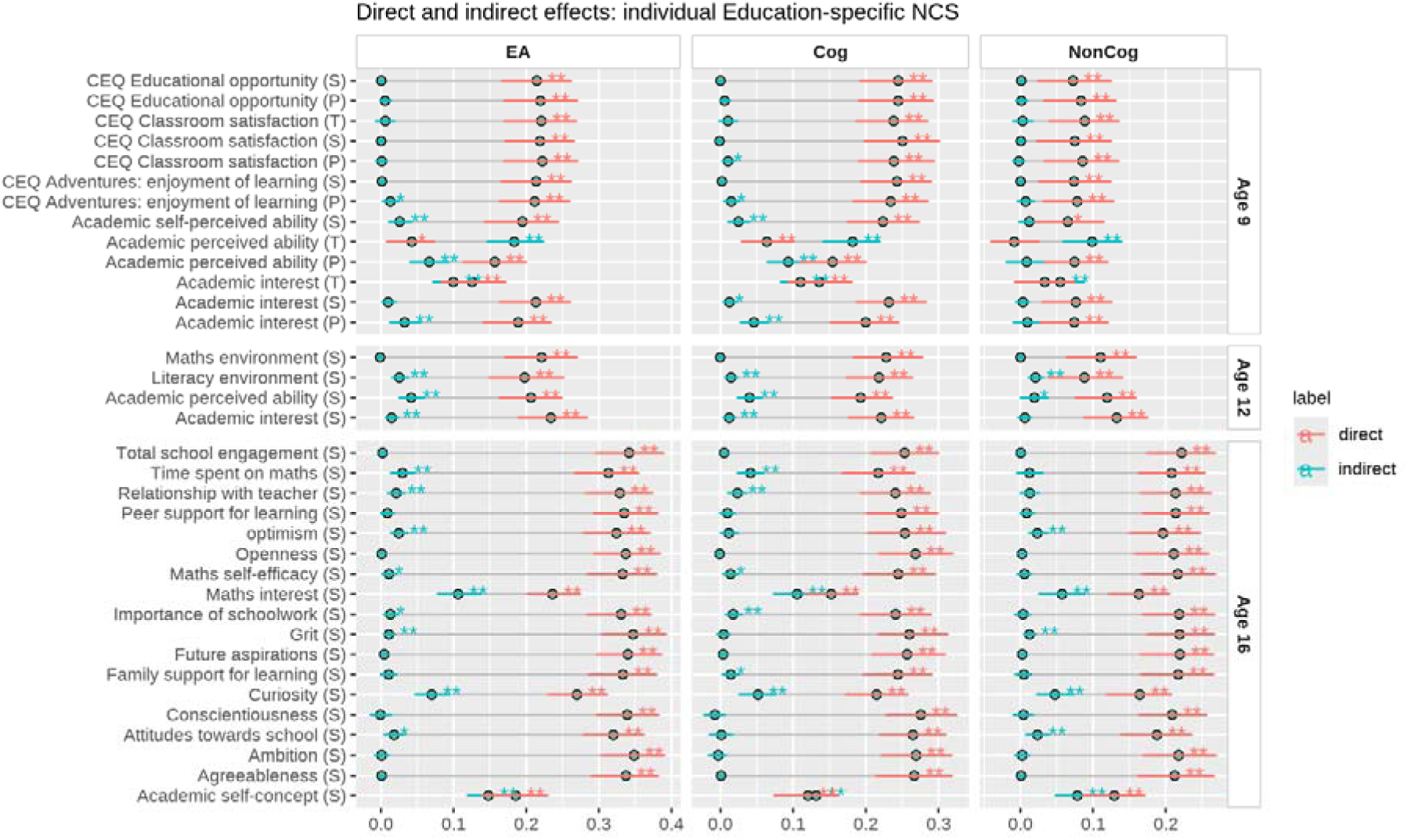
Direct and indirect effects of polygenic score associations with academic achievement as mediated by individual education-specific non-cognitive skills (NCS). Variables labelled with (P), (T), and (S) represent different informants: (P) = Parent-rated; (T) = Teacher-rated; (S) = Self-rated. Each dot represents the effect size (standardized ß coefficient) for direct (red) and indirect (blue) polygenic score effects on academic achievement at ages 9, 12 and 16. EA = educational attainment polygenic score^14^ (left panel), Cog = cognitive polygenic score^29^ (middle panel), and NonCog = non-cognitive polygenic score^29^ (right panel). Error bars around each dot indicate standard errors. * = p < 0.05, ** = p < 0.01 after applying FDR correction.

Although for self-rated non-cognitive measures, mediation effects increased developmentally, for example, for academic self-perceived ability and academic interest (**Fig 2**).

By the end of compulsory education, the mediating role of self-rated non-cognitive measures was substantial. For example, the indirect (mediated) effects of academic self-concept and maths interests measured at age 16 accounted for 45% and 32% of the association between the EA polygenic score and academic achievement at age 16, respectively (**left panel of Fig 2**). Moreover, when we partitioned polygenic score effects into cognitive and non-cognitive genetics, we observed that academic self-concept and interest mediated both polygenic score associations with academic achievement. For the Cog polygenic score-academic achievement association, mediation effects accounted for 52% and 42% of the association, respectively (**middle panel of** **Fig** 2); while for the NonCog polygenic score for 38% and 27%, respectively (**right panel of Fig 2**). Overall, mediation effects observed for individual non-cognitive measures were highly consistent across the three polygenic scores considered (**Fig 2 and Supplementary Tables 7-9)**.

### Non-cognitive skills mediate genetic disposition of academic achievement longitudinally

Given the consistent mediation effects of education-specific non-cognitive skills that emerged from our analyses at the same developmental stage, we next examined time-lagged effects longitudinally, such as whether non-cognitive skills measured at age 9 mediated polygenic score effects on academic achievement at ages 12 and 16. We observed persistent mediation effects even in these time-lagged analyses, although effects were slightly attenuated in some, but not all, cases (**Fig 3**). For example, the role of self-rated non-cognitive skills at age 12 as mediator of the association between EA polygenic score and academic achievement at age 16 remained substantial (**right panel of Fig 3**). Findings were highly consistent when we examined Cog and NonCog polygenic score effects (**see middle and bottom panel of Fig 3 and Supplementary Table 10)** and for individual non-cognitive measures (**see Supplementary Table 11**).

**Fig 3:**
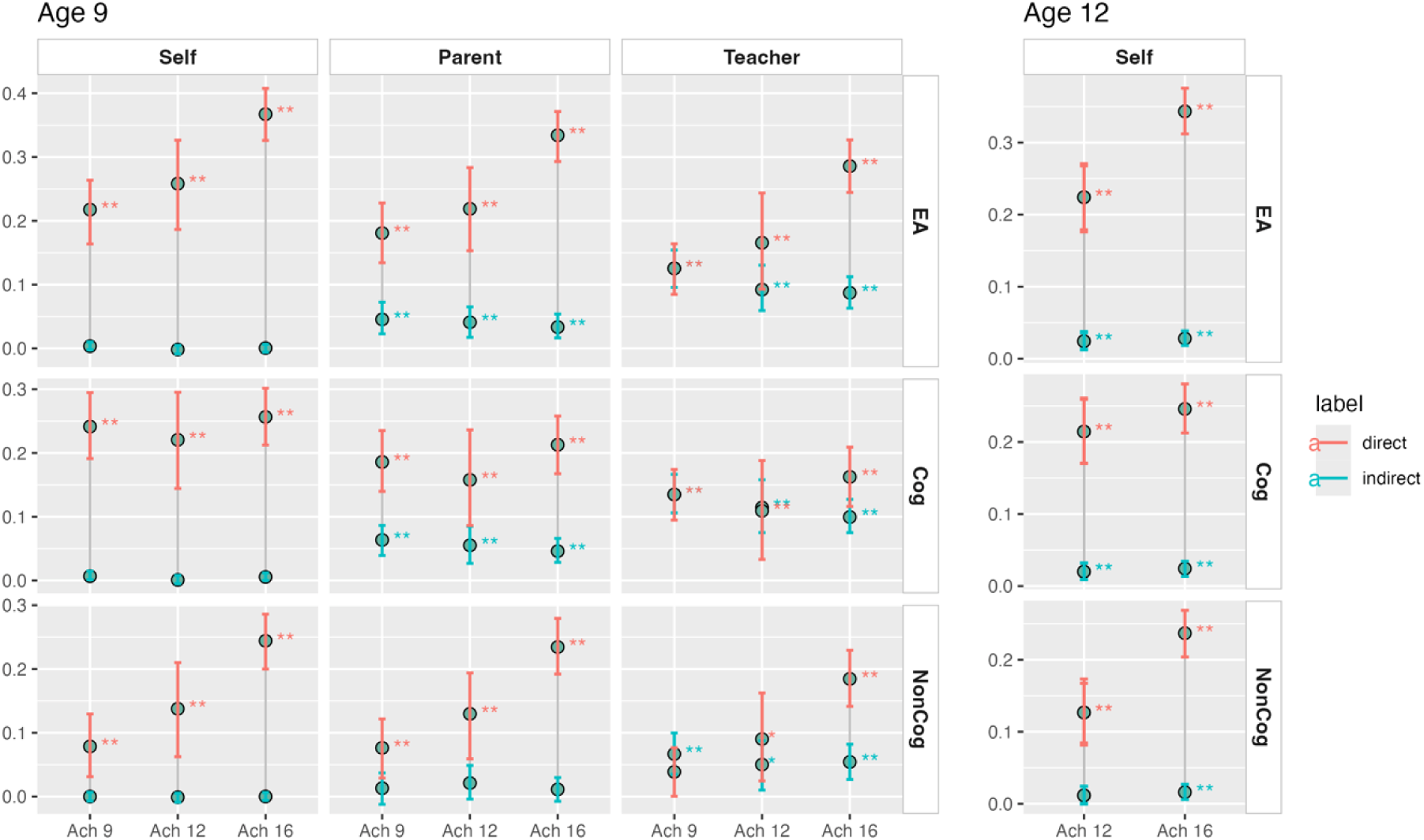
Time-lagged mediating effects of Education-specific non-cognitive skills at age 9 (left panel) and age 12 (right panel) on the polygenic score (PGS) predictions of academic achievement over development. The colour represents the direct (red) and indirect (blue) effects (standardised beta coefficient) of the educational attainment^14^ (EA; top panel), cognitive^29^ (Cog) and non-cognitive^29^ (NonCog; bottom panel) polygenic scores on academic achievement at ages 9, 12, and 16. Error bars indicate the standard error around the effect size. * = p < 0.05, ** = p < 0.01 after FDR correction.

### Within-family mediation analyses

We applied multi-level mediation models (see **Methods**) to separate polygenic score associations into within- and between-family effects and further disentangle the processes underlying the role of non-cognitive skills in these associations. The number of sibling (DZ) pairs included in the analyses ranged between 1,104 to 2,895, depending on the data collection wave and measures included. Family-level intraclass correlations (ICCs) for latent non-cognitive factors ranged between 0.23 and 0.52 (see **Supplementary Table 12**), indicating moderate to substantial family-related effects on the development of non-cognitive skills.

Consistent with previous research^13,29,35^, we found that, the associations between EA polygenic score and academic achievement were attenuated when looking at differences within families. However, young people’s non-cognitive skills remained significant mediators of the association between the educational attainment polygenic score and academic achievement even when comparing siblings (**Fig 4 top panel and Supplementary Table 13**). Siblings with a higher education-related polygenic score showed on average greater academic achievement, and this association was partly accounted for by higher levels of non-cognitive skills. Notably, our finding of a developmental increase in the mediating role of self-reported non-cognitive skills replicated at the within-family level (**see left panel of Fig 5**).

**Fig. 4:**
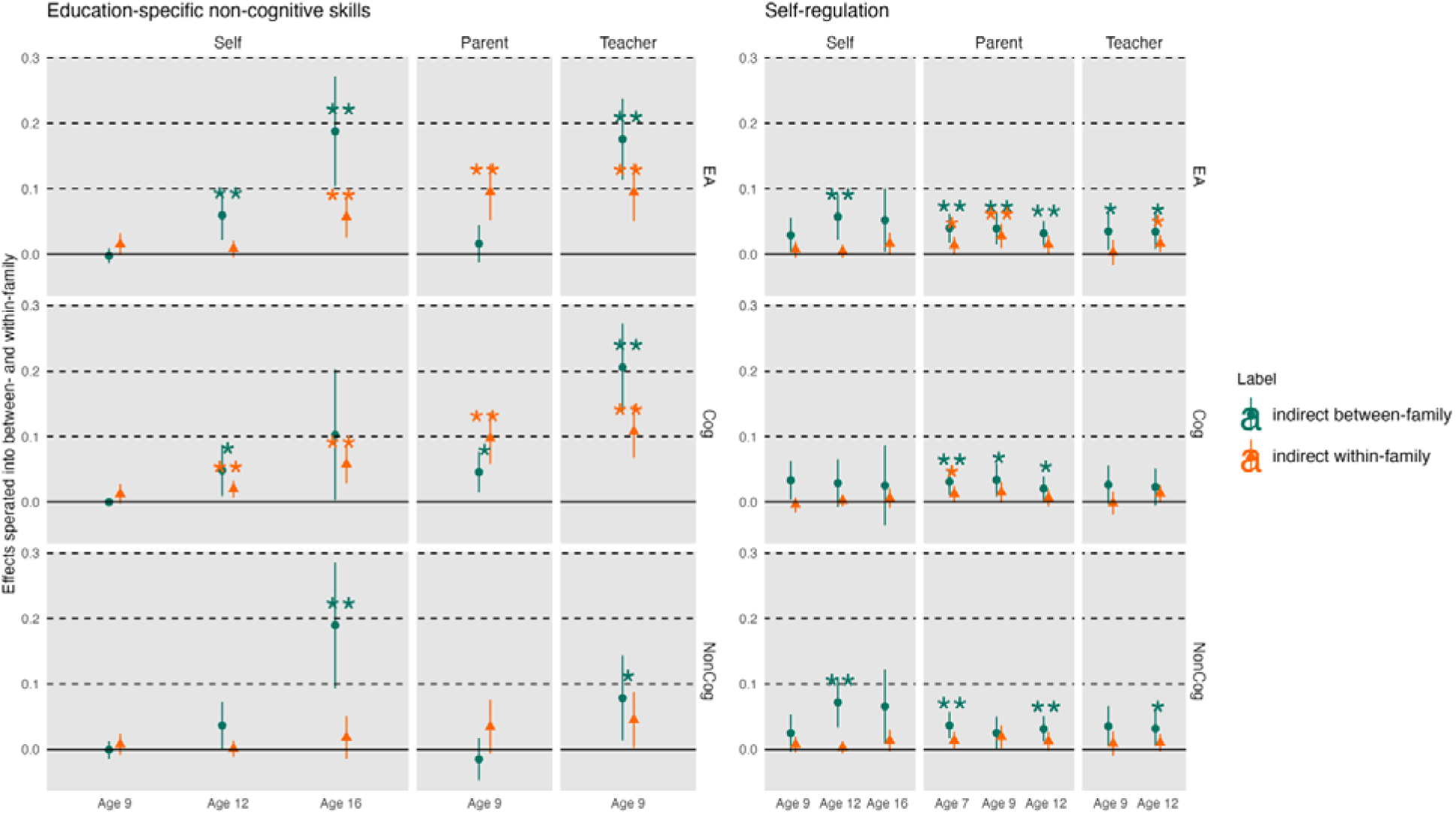
Mediation effects of the non-cognitive latent factors in the polygenic score (PGS) predictions of academic achievement separated into between- and within-family effects. The dots represent the effects of the PGS predictions (standardised beta coefficient) of academic achievement at ages 7, 9, 12 and 16 partitioned into between- and within-family effects. Education-specific non-cognitive skills: left panel. Self-regulation: right panel. Educational attainment polygenic score (EA PGS)^14^: top panel. Cognitive polygenic score (Cog PGS)^29^: middle panel. Non-cognitive polygenic score (NonCog PGS)^29^: bottom panel. * = p < 0.05, ** = p < 0.01 after applying FDR correction.

For parent-rated non-cognitive skills, mediation effects were mainly evident at the within-family level rather than between-families (i.e., between-family mediation effect: ß = 0.02 [95% CI, -0.01, 0.05]; within-family mediation effect: ß = 0.10 [95% CI, 0.05, 0.14]), accounting for 83% of the total mediation effects. Polygenic score associations with academic achievement that are unique to each child, rather than shared by siblings, were more likely to be mediated parents’ perceptions of non-cognitive profiles unique to each child. While this was observed for education-specific non-cognitive skills, associations were different for self-regulation. When considering children’s unique self-regulation profiles, most mediation effects were attenuated and did not reach significance in within-family analyses. We observed a similar pattern of results when repeating mediation analyses separating education-related genetics into cognitive and non-cognitive polygenic scores (**middle and bottom panel of Fig 4; Supplementary Tables 14 and 15**). Results were also consistent when we examined the mediating role of each education-specific non-cognitive skills individually (**Supplementary Table 16**).

We further investigated whether within-family mediation effects could be observed longitudinally. We found that time-lagged mediation effects for education-specific non-cognitive measures persisted even when looking within families. For example, parent-rated education-specific non-cognitive skills measured at age 9 mediated EA polygenic score associations with academic achievement at both age 12 (ß = 0.08 [95% CI, 0.03, 0.12]) and age 16 (ß = 0.08 [95% CI, 0.05, 0.11]). The effect size was comparable to the effect size observed cross-sectionally in the association between the EA polygenic score and academic achievement measured at age 9 (that is, ß = 0.10 [95% CI, 0.05, 0.14]). The full results are presented in **Supplementary Tables 17-19** and provide evidence for the long-lasting mediating role of education-specific non-cognitive skills in the association between polygenic score and achievement across compulsory education. When examining the effects of each education-related non-cognitive skill individually, we found that parent-rated academic perceived ability and academic interest evinced the strongest mediation effects (ß = 0.09 [95% CI, 0.04, 0.15] and ß = 0.08 [95% CI, 0.05, 0.11], see **Supplementary Table 20)**.

### Adjusting for general cognitive ability (g) and family socio-economic status (SES)

Since previous research has shown that students’ non-cognitive skills correlate with students’ cognitive ability^26,29^, we tested the possibility that cognitive ability could account for the mediation effects we observe. Specifically, we repeated our mediation analyses, including general cognitive ability as a second mediator (see **Methods**). Results showed that non-cognitive skills remained significant, although slightly attenuated, mediators of the polygenic scores-achievement associations (**Supplementary Figure 3 and Supplementary Table 21**). Results were similar across compulsory education and across the different polygenic scores (**Supplementary Figures 4 and 5; Supplementary Table 22**). Non-cognitive skills also remained significant mediators after accounting for family socio-economic status (**Supplementary Figure 6 and Supplementary Table 23**), with similar results observed across polygenic scores and compulsory education (**Supplementary Figures 7 and 8; Supplementary Table 22**).

## Discussion

While it is well-established that genetic differences between students partly contribute to their differences in academic achievement^6,7^, how genetic differences come to manifest as differences in academic achievement remains under-investigated. To address this major gap in our knowledge, the present study provides an in-depth examination of the role that *non-cognitive skills* (e.g., academic motivation, attitudes and goals) play in the association between genetic disposition and observed academic achievement across compulsory education. Our analyses revealed three key findings. First, non-cognitive skills, particularly those skills closely linked to learning (e.g., academic interest, curiosity, and academic self-perceived ability), play a critical role in translating education-associated genetic dispositions into observed academic achievement. Second, our time-lagged analyses showed that the mediating role of non-cognitive skills persisted across development. Third, these mediation effects could also be observed when examining differences between siblings. Mediation effects were small yet consistent and robust to adjusting for family socio-economic status and general cognitive ability, highlighting the independent contribution of non-cognitive skills to academic success.

Our cross-sectional analyses showed that the latent dimensions of education-specific non-cognitive skills and self-regulation both mediated the associations between education-related genetic disposition and academic performance. However, while the role of self-regulation was stable across compulsory education, the mediating role of education-specific non-cognitive skills increased developmentally, pointing to the growing importance of students’ perceived non-cognitive profiles and experiences in their academic journeys. This developmental increase is consistent with the possibility that, as they grow up, children become more aware of their aptitudes and appetites towards learning. As they gain greater self-awareness and autonomy, students might become increasingly more able to shape their environmental contexts in ways that allow them to cultivate these non-cognitive skills and, in turn, foster their academic performance^36^.

The magnitude of mediation effects also differed depending on raters. Teacher reports of education-specific non-cognitive skills evinced the strongest effects, followed by parent reports, and self-reports. These stronger effects may reflect rater bias, as teachers evaluated both students’ performance and their learning-related attitudes^37^. However, they may also reflect the greater expertise and objectivity that characterise teachers’ assessments^38^. This is supported by our finding that teacher-rated non-cognitive skills had long-lasting mediation effects in our time-lagged analyses, since different teachers assessed achievement at different ages. Teachers have first-hand experience of how children express their attitudes and appetites within academic settings and might, therefore, be able to capture students’ non-cognitive competencies related to learning more accurately^39^.

With a set of fine-grained analyses, we investigated which aspects of non-cognitive skills played a greater role in mediating the link between genetic disposition and academic achievement across development. First, we observed that student-centred academic attitudes and perceptions, such as academic interest, curiosity, and perceived ability, evinced stronger mediation effects as compared to perceptions of the learning environment that were external to each individual, such as perception of the classroom or the learning environments. These findings align with previous work, which emphasised the critical role of intrinsic, student-cantered non-cognitive skills (e.g., intellectual curiosity and academic self-concept) in gene-environment transactions^36,40^, driving students to actively engage in academic settings beyond what environmental factors alone can achieve^41^. This might be particularly useful knowledge for interventions aimed at fostering learning. Interventions aimed at enhancing students’ academic self-concept, interests, and motivations, such as inquiry-based learning^42^, positive feedback^43^, and goal-setting^44^, might be particularly effective.

We examined whether the mediating role of non-cognitive skills was more strongly linked to cognitive or non-cognitive genetics. We found that mediation effects were comparable when separating education-associated genetic effects into cognitive and non-cognitive genetics.

Interestingly, for some student-centered non-cognitive skills such as academic interest and academic self-concept, mediating effects were stronger for the path from cognitive genetics to academic achievement. This is consistent with observations of moderate associations between cognitive and non-cognitive skills across compulsory education^26,29^. In fact, although the term *non-cognitive* skills was coined to highlight their independent contribution to academic performance beyond cognitive skills, the two constructs are related and developmentally intertwined. These results also align with propositions that non-cognitive skills like interest, curiosity and self-discipline help students leverage their cognitive competencies more effectively, therefore facilitating academic performance^45^

Our time-lagged analysis confirmed that the mediating role of non-cognitive skills lasted throughout compulsory education. These findings reinforce the pivotal role of non-cognitive skills in translating genetic dispositions into academic achievement, and underscore the importance of fostering non-cognitive skills to enhance students’ long term academic outcomes, consistent with previous studies on the effectiveness of interventions for improving non-cognitive skills. For instance, interventions focusing on self-control and social skills at age 7 were found to have a positive impact on academic achievement in early adulthood^46^. A recent systematic review and meta-analysis showed that early-life interventions aimed at improving self-regulation and perseverance were associated with modest improvements in later-life academic and psychosocial outcomes^47^.

Our within-family analyses provided us with a more nuanced understanding of the potential mechanisms through which genetic dispositions and non-cognitive experiential processes combine and contribute to differences between students in academic achievement. Examining differences between siblings, dizygotic twins in this case, we were able to separate the effects of family-wide processes from those processes that are unique to each child within a family. We found that the mediating role of non-cognitive skills could be observed not only between-families, but also when looking at differences between siblings, accounting for a substantial portion of the total effect. Each children’s unique genetic dispositions were associated with academic achievement partly through their distinct non-cognitive profiles beyond those genetic, environmental and demographic factors that are common to family members, which include assortative mating^34^, population stratification^33^, and passive gene-environment correlation. This underscores the key role played by individual-specific non-cognitive profiles in propelling children down different learning pathways. Our findings are in line with transactional models of human development rooted in evocative/active gene-environment correlation^36,48^ that proposes that, partly in line with their genetic dispositions, children actively seek out different environmental experiences on the basis of their non-cognitive characteristics (e.g., interests, preferences, and aptitudes), and in turn, these will lead to different learning outcomes.

A number of limitations should be acknowledged. First, the reliance on polygenic scores as proxies for genetic influences on academic achievement is an inherent limitation, as these scores are imperfect representations of the complex genetic architecture underlying academic performance^14,49^. This is particularly relevant to our non-cognitive polygenic score analyses, since this score is indirectly constructed by isolating those genetic variants related to education that are not associated with cognitive skills, and not directly from a GWAS of actual non-cognitive measures^29^. It would be of interest to replicate these results when adequately powered GWAS of non-cognitive skills will become available. Second, our non-cognitive measures may not fully capture the breadth and depth of factors influencing individuals’ non-cognitive profiles relevant to academic success ^23,50^. Extant literature on the links between non-cognitive skills and educational outcomes is complex and characterised by inconsistencies, low-quality studies and publication bias^47^. Greater clarity regarding the taxonomy of non-cognitive skills that relate to education, better measures, and high-quality longitudinal studies are urgently needed.

Third, we cannot exclude potential genetic confounding in our mediation analyses. Mediation pathways may be under-corrected for genetic influences in phenotypic variables^51–53^, which could lead to an overestimation of the genetic effects mediated by non-cognitive skills. Fourth, adjusting for heritable covariates such as family SES and general cognitive ability, could introduce collider bias, potentially distorting the relationship between non-cognitive skills and academic achievement^54^. Fifth, our study was conducted within a UK-based twin sample, primarily focused on individuals of White-European ancestry, which, although representative of the UK population for their birth cohort, limits the generalisability of our findings to other populations with different socio-cultural and ancestral backgrounds. Increasingly diverse samples and methodological advances in genetic research, such as multi-ancestry GWASs^55^ and novel GWAS methods^56^, will help bridge this gap in future studies.

Lastly, with the emergence of new methodologies to differentiate between-family and within-family genetic effects, we plan to extend our research and continue triangulating evidence using multiple approaches^57^.

## Methods

### Participants

Our study involved twins who were part of the Twins Early Development Study (TEDS)^58^. TEDS has gathered information from twins born in England and Wales between 1994 and 1996, along with their parents, at various intervals spanning childhood, adolescence, and early adulthood, commencing from birth. The initial data collection involved the participation of over 13,000 twin pairs, and after almost three decades, TEDS maintains an active membership of over 10,000 families. TEDS sample remains largely representative of the demographic characteristics of the UK population for their respective generation in terms of ethnicity and socio-economic status^59,60^. The subsample included in our current analyses comprised 5,016 individuals (2,895 DZ sibling pairs for the within-family design) whose families had contributed data on academic achievement, non-cognitive skills, and had available genotype information. The gender distribution within the sample was approximately 51.1% female and 48.9% male. Our examination spanned four waves of data collection: age 7 (Mean age = 7.20), age 9 (Mean age = 9.03), age 12 (Mean age = 11.52), and age 16 (Mean age = 16.31). Mean ages are derived from teacher assessment of academic achievement. Participants with significant medical, genetic, or neurodevelopmental conditions were excluded from our analyses, resulting in a variable sample size ranging from 1,516 to 5,016 due to the inclusion of distinct variables.

### Measures

Data were collected by means of questionnaires and tests administered to parents, teachers, and the twins themselves by post, telephone, and online, as described in detail in this overview of the TEDS study^60^ and in the TEDS data dictionary (https://www.teds.ac.uk/datadictionary/home.htm).

#### Academic achievement

At ages 7, 9 and 12, academic achievements were provided by the teachers who assessed students’ performance based on the UK National Curriculum guidelines designed by the National Foundation for Educational Research (NFER; http://www.nfer.ac.uk/index.cfm) and the Qualifications and Curriculum Authority (QCA; http://www.qca.org.uk). At age 7, academic achievement was measured as a composite of standardised academic performance across two subjects: English and mathematics. At age 9 and 12, a composite score of standardised academic performance across three subjects: English, mathematics, and science.

Academic achievement at age 16 was measured as the mean grade score for the General Certificate of Secondary Education (GCSE) three subjects (English, mathematics and science). GCSEs are standardized tests taken at age 16, which in the UK for the TEDS sample was the end of compulsory education. The exams are graded on a scale ranging from A* to G, with a U grade assigned for unsuccessful attempts. The grades were coded on a scale from 11 (A*) to 4 (G, the lowest passing grade), and the mean of the grade obtained from the GCSE core subjects was used as our measure of academic achievement at age 16. Data on GCSE exam results were collected from parental and self-reports via questionaries sent over mail or telephone. Our previous research has shown that teacher ratings and self-reported GCSE grades are valid, reliable, and correlate very strongly with standardized exam scores taken at specific moments in the educational curriculum (Key Stages) obtained from the National Pupil Database^61^.

#### Non-cognitive skills

Below we provide a brief description of all the measures included in the current study. More details can be found in the TEDS data dictionary (https://www.teds.ac.uk/datadictionary), which provided detailed description of each measure and information on the items included in each construct.

### Education-specific non-cognitive skills

At **age 9** data on education-specific non-cognitive skills were collected from twins themselves, their parents and teachers. Measures of academic self-perceived ability^62^, academic interest^62^ and the Classroom Environment Questionnaire (CEQ^63^) were available from all raters. The CEQ included the following subscales rated by parents and twins: (1) CEQ classroom satisfaction scale; (2) CEQ educational opportunities scale; (3) CEQ adventures scales, assessing enjoyment of learning. Ratings on the CEQ classroom satisfaction scale were also provided by the teachers.

At **age 12** data on education-specific non-cognitive skills were collected from self-reports, parents and teachers. We collected the following measures: academic interest^62^, academic self-perceived ability^62^, the literacy environment questionnaire^64,65^ and the mathematics environment questionnaire^66^. The questionnaires asked several questions related to literacy and mathematics, for example: *Reading is one of my favourite activities; When I read books, I learn a lot*; and *In school, how often do you do maths problems from text books?* all rated on a four-point Likert scale.

At **age 16** education-specific non-cognitive skills were assessed via self-reports provided by the twins. The battery of education-specific non-cognitive constructs included the following measures: **Academic self-concept**: 10 items of a brief academic self-concept scale (adapted from^67^, e.g., ‘*I like having difficult work to do* and *I am clever’)* rated on a 5-point Likert scale.

**School engagement**^68^: a composite includes 5 subscales: future aspirations and goals; control and relevance of schoolwork; teacher-student relations; peer support for learning; family support for learning. The scale includes items rated on four-point Likert scale, such as: ‘*School is important for achieving my future goals’*, ‘*I feel like I have a say about what happens to me at school’*, ‘*I enjoy talking to the teachers at my school’*, and ‘*When I have problems at school, my family/carer(s) are willing to help me’*).

**Grit** was measured with 8 items from the Short Grit Scale (GRIT-S)^69^ asking the twins to report on their academic perseverance answering questions (e.g., ‘*Setbacks don’t discourage me*, and *I am a hard worker’)* rated on a 5-point Likert scale.

**Academic ambition**^70^ was assessed with 5 items asking participants to rate statements like the following: ‘*I am ambitious* and *achieving something of lasting importance is the highest goal in life’*, on a 5-point Likert scale.

**Time spent on mathematics** was measured with 3 items asking participants how much time every week they spent in, including ‘*Regular lessons in mathematics at school’, ‘Out-of school-time lessons in mathematics’*, and ‘*Study or homework in mathematics by themselves’*.

**Mathematics self-efficacy** (OECD Programme for International Student Assessment (PISA)) was assessed with 8 items rated on four-point Likert scale. The scale including questions to measure how confident the student felt about having to conduct different mathematics tasks (e.g., ‘*Calculating how many square metres of tiles you need to cover a floor* and *Understanding graphs presented in newspapers’*).

**Mathematics interest** (OECD Programme for International Student Assessment (PISA)) was assessed with 3 questions, asking participants about how their interest in mathematics, such as: ‘*I do mathematics because I enjoy it* and *I am interested in the things I learn in mathematics’*.

**Curiosity** was assessed with 7 items^71^ rated on 7-point Likert scale. Students were asked to rate statements including: ‘*When I am actively interested in something, it takes a great deal to interrupt me* and *Everywhere I go’, ‘I am looking out for new things or experiences’*.

**Attitudes towards school** was assessed using the OECD Programme for International Student Assessment (PISA) attitudes to school measure which included 4 items rated on a 4-point Likert scale, such as: ‘*School has helped give me confidence to make decisions’* and ‘*School has taught me things which could be useful in a job’*.

### Self-regulation

The Strengths and Difficulties Questionnaire (SDQ)^72^ scale was used to evaluate emotional and behavioural self-regulation across all ages. Data on domain-general self-regulation abilities was collected from parents, teachers, and self-reports of the twins. The SDQ comprises five subscales: conduct problems, peer problems, hyperactivity, emotional difficulties, and prosocial behaviour. Scores from all subscales, except prosocial behaviour, were reversed to ensure higher scores indicate better self-regulation skills. Parents rated self-regulation skills at age 7, while assessments at ages 9 and 12 involved inputs from parents, teachers, and the twins. At age 16, twins reported their own self-regulation skills.

#### Cognitive ability

Cognitive ability at **age 7** was evaluated through four tests administered via telephone by trained research assistants. Verbal cognitive ability was measured using two tests adapted from the Wechsler Intelligence Scale for Children (WISC-III)^73^: a 13-item Similarity test and an 18-item Vocabulary test. Nonverbal cognitive ability was evaluated with a 9-item Conceptual Groupings Test^74^ and a 21-item WISC Picture Completion Test^73^. Composite scores for verbal and nonverbal abilities were calculated by taking the average of the standardized scores within each respective domain. Furthermore, a general cognitive ability composite (*g*) was established by averaging the standardized scores obtained from the two verbal and two nonverbal tests.

Cognitive ability at **age 9** was evaluated through four booklets sent to TEDS families by mail. Verbal skills were assessed using the initial sections (20 items) of the WISC-III-PI Words and General Knowledge tests^75^. Nonverbal abilities were measured with the Shapes test (CAT3 Figure Classification^76^ and the Puzzle test (CAT3 Figure Analogies)^76^. Composite scores for verbal and nonverbal skills were created by averaging the standardized scores within each area. Additionally, a *g* composite was calculated by averaging the standardized scores from the two verbal and two nonverbal tests.

Cognitive ability at **age 12** was evaluated through four tests administered online. Verbal skills were assessed using the complete versions of the verbal tests conducted at age 9: the entire 30 items from the WISC-III-PI Words test^75^ and 30 items from the WISC-III-PI General Knowledge test^75^.

Nonverbal abilities were measured with the 24-item Pattern test (derived from Raven’s Standard Progressive Matrices)^77^ and the 30-item Picture Completion test (WISC-III-UK)^73^. Composite scores for verbal and nonverbal abilities were derived by averaging the standardized scores within each respective domain. Furthermore, a *g* composite was computed by averaging the standardized scores from the two verbal and two nonverbal tests.

Cognitive ability at **age 16** was assessed through a comprehensive evaluation consisting of one verbal test and one nonverbal test administered online. Verbal proficiency was measured using a modified iteration of the Mill Hill Vocabulary test^78^, while nonverbal abilities were gauged through an adapted version of the Raven’s Standard Progressive Matrices test. A *g* composite was determined by averaging the standardized scores derived from the results of both tests.

#### Family socio-economic status

We used socioeconomic status data collected at the first contact for the models at ages 7, 9, and 12. For the models at age 16, we used the socioeconomic status data available at that age. Specifically, at first contact, TEDS twins’ parents were contacted via postal mail and asked to complete a questionnaire concerning their educational background, employment status, and the age at which the mothers had their first child. A composite measure of socioeconomic status was created by standardizing these three variables and calculating their average. At the age of 16, socioeconomic status information was collected through a web-based questionnaire, and a comprehensive score was generated by averaging the standardized values of five items: household income, the highest educational attainment of both parents, and the employment status of both parents.

#### Genetic data

Two different genotyping platforms were utilized due to genotyping being conducted in two distinct phases, with a 5-year interval between them. Affymetrix GeneChip 6.0 SNP arrays were employed to genotype 3,665 individuals, while Illumina HumanOmniExpressExome-8v1.2 arrays were used for genotyping 8,122 individuals (including 3,607 DZ co-twin samples). Following quality control procedures, genotypes from a total of 10,346 samples (comprising 3,320 DZ twin pairs and 7,026 unrelated individuals) were deemed acceptable, with 3,057 individuals genotyped on Affymetrix and 7,289 individuals genotyped on Illumina arrays. The final dataset included 7,363,646 genotyped or well-imputed SNPs. For further details regarding the handling of these samples, refer to the provided source^79^.

#### Polygenic score

Following the implementation of DNA quality control measures recommended for chip-based genomic data, we developed genome-wide polygenic scores (PGS) utilizing summary statistics obtained from four distinct genome-wide association studies: *educational attainment*^14^, *cognitive ability*^29,80^ and *non-cognitive skills*^29,31^. Each PGS was computed as the weighted sum of an individual’s genotype across all single nucleotide polymorphisms (SNPs), with adjustments made for linkage disequilibrium using LDpred^81^. LDpred is a Bayesian shrinkage method that adjusts for local linkage disequilibrium (the correlation between SNPs) by utilizing data from a reference panel. In our study, we used the target sample (TEDS), limited to unrelated individuals, along with a prior reflecting the genetic architecture of the trait. We constructed polygenic scores (PGS) using an infinitesimal prior, assuming that all SNPs contribute to the genetic architecture of the trait. This approach has been shown to work well for highly polygenic traits like educational attainment (EA). In our regression analyses, both the cognitive (Cog) and non-cognitive (NonCog) PGSs were included in multiple regression models, along with covariates such as age, sex, the first ten principal components of ancestry, and genotyping chip and batch. To address the non-independence of observations, we used the generalized estimating equation (GEE) approach^82^ from the R package.

### Analytic strategies

All scripts will be made available on our research lab GitHub page (https://github.com/CoDEresearchlab).

#### Data preparation

To ensure thorough control for potential confounding factors and to effectively evaluate the mediating impact of these measures on the association between polygenic scores and academic performance, all non-cognitive skill measures and academic achievements underwent regression against age and sex effects based on the information on the same measure.

#### Construction of latent factors of non-cognitive skills and general cognitive ability

Confirmatory factor analysis (CFA)^83^ was utilised to establish latent constructs representing general cognitive ability and non-cognitive skills across various developmental stages. In alignment with established research on general cognitive ability (*g*) and prior investigations within the TEDS dataset, we formed a single factor for *g* at each developmental phase. These *g* factors were constructed by aggregating the weighted loadings of two verbal and two nonverbal tests, as outlined in the **Measures** section and **Supplementary Table 23**.

CFA was also utilised to create dimensions reflecting non-cognitive skills. Drawing from previous meta-analytic research on the non-cognitive factors influencing educational outcomes, we adopted a theoretical differentiation between education-specific non-cognitive traits (such as academic related attitudes, motivation, and goals) and broader, more universally applicable measures of self-regulation (encompassing behavioural and emotional regulation). Consequently, distinct factors were established for a) education-specific non-cognitive skills (NCS) and b) domain-general self-regulation skills, considering the various measures available at each age for each evaluator. Please refer to **Supplementary Table 24** and **25** for the factor loadings and model fit indices associated with these constructs.

#### Mediation analyses

Following the removal of outliers (scores outside +/- 4 standard deviations), we employed Structural Equation Modelling (SEM) for conducting mediation analyses^84^ using *lavaan* in R^85^. A comprehensive description of the mediation models is provided in **Supplementary Note 2**. These mediation models enabled us to dissect the effect size of each polygenic score’s prediction on academic achievement throughout development into direct and indirect effects mediated by both latent dimensions of, and individual non-cognitive skill measures. To address the risk of multiple testing, we employed the Benjamini-Hochberg false discovery rate correction (FDR)^86^. Furthermore, to address the non-independence of observations in the sample (i.e., relatedness), we randomly selected one twin from each pair for analysis. Also, we applied bootstrap method^87^ within *lavaan* package, which allows us to assess the variability of our estimates without relying on the assumption of normality in the sampling distribution.

#### Cross-sectional analyses

We first conducted mediation models for the non-cognitive skills and academic achievement at the same developmental stage across compulsory education, examining the pathway from polygenic scores to non-cognitive skills and academic achievement measured at the same age. For example, we investigated whether the education-specific non-cognitive skills assessed at age 9 mediated the polygenic score effects on academic achievement at the same age. We selected mediators based on data availability during the same collection wave as each academic achievement outcome.

#### Longitudinal mediation analyses

We further applied longitudinal models, using non-cognitive skills measured at earlier ages as mediators for later academic achievement to investigate whether the non-cognitive skills mediated genetic disposition with academic achievement longitudinally. For instance, we conducted mediation models by including education-specific non-cognitive skills assessed at age 9 and academic achievement at ages 12 and 16.

#### Two-mediator mediation analyses

To account for potential confounding effects, we utilized *two-mediator mediation models*^88^ to expand our examination of mediation effects and explore whether the mediating role of non-cognitive skills was primarily influenced by either family socioeconomic status (SES) or general cognitive ability (*g*). Therefore, we reiterated our mediation models by simultaneously considering each non-cognitive skill alongside family SES and the *g* composite.

#### Multi-level mediation analysis: separating between from within-family effects

We further utilized *1-1-1 two-level mediation models*^89^ to disentangle within-family effects from between-family effects. This statistical framework enabled us to assess the indirect impact of a predictor on an outcome while considering clustered mediation data by mean-centring the siblings’ measurements and taking the departure from the mean for each sibling. In our analyses, we organized our data by family, with each family representing a cluster comprising two members (the two dizygotic twins). By employing *1-1-1* multilevel mediation models, we could separate the between-sibling polygenic score effects, as well as mediation effects. Between-family effects estimated from samples of unrelated individuals are typically biased by sibling-invariant family level demographic factors such as genetic nurture and population stratification and passive gene-environment correlation ^17,19,90^. Within-family analyses rely on the randomization of allele transmission during meiosis, ensuring that siblings have an equal chance of inheriting any given allele, independent of environmental processes. Consequently, genetic differences between siblings are unaffected by environmental factors making within-family effects robust to major sources of environmental contributions. Our within-family mediation estimates are therefore indicative of how each sibling’s genetic disposition towards education combines with individual-specific socio-emotional processes to lead to differential academic outcomes through evocative/active gene-environment correlation processes.

In these analyses, only dizygotic (DZ) twins were included, as our objective was to explore how differences in polygenic scores between siblings predicted variation in academic achievement through disparities in the non-cognitive skills.

## Code availability statement

The code is available at: https://github.com/CoDEresearchlab

## Acknowledgements

This study was funded by a starting grant awarded to MM by the School of Biological and Behavioural Sciences at Queen Mary, University of London. M.M. is supported by a Jacobs Foundation Research Fellowship. QZ is funded by the QMUL-Chinese Scholarship Council joint PhD Scholarship. TEDS is supported by a programme grant to RP from the UK Medical Research Council (MR/V012878/1 and previously MR/M021475/1), with additional support from the US National Institutes of Health (AG046938).

## Author contributions

Conceived and designed the study: M.M., Q.Z. Analysed the data: Q.Z. Supervised the work: M.M.. Wrote the paper: Q.Z., M.M., with helpful contributions from A.A., J.W., T.T.M., L.R., K.R., and R.P.. All authors contributed to the interpretation of data, provided feedback on drafts, and approved the final draft.

