## Supplementary Information for "From genetic disposition to academic achievement: The mediating role of non-cognitive skills across development"

### Table of Contents

*Supplementary Notes* 3

**Supplementary Note 1: Descriptive statistics.** 3

**Supplementary Note 2: Description of mediation models.** 3

*Supplementary Figures* 6

**Supplementary Figure 1: Correlation matrix between PGS and academic achievement.** 6

**Supplementary Figure 2: Correlations between latent factors of non-cognitive skills, academic achievement and education and cognition-associated polygenic scores.** 7

**Supplementary Figure 3: Comparison of indirect effects of educational attainment prediction between g-uncorrected/ g-corrected noncognitive skills using two-mediators model.** 8

**Supplementary Figure 4: Comparison of indirect effects of cognitive prediction between g-uncorrected/ g-corrected noncognitive skills using two-mediators model.** 9

**Supplementary Figure 5: Comparison of indirect effects of noncognitive prediction between g-uncorrected/ g-corrected noncognitive skills using two-mediators model.** 10

**Supplementary Figure 6: Comparison of indirect effects of educational attainment prediction between SES-uncorrected/ SES-corrected noncognitive skills using two-mediators model.** 11

**Supplementary Figure 7: Comparison of indirect effects of cognitive prediction between SES-uncorrected/ SES-corrected noncognitive skills using two-mediators model.** 12

**Supplementary Figure 8: Comparison of indirect effects of noncognitive prediction between SES-uncorrected/ SES-corrected noncognitive skills using two-mediators model.** 13

### Supplementary Notes

#### Supplementary Note 1: Descriptive statistics.

We provide descriptive statistics for the variables used in present study, which includes various measures of noncognitive skills (both individual and latent factors), overall academic achievement and polygenic scores collected from different raters at different ages. Only genotyped participants were included and sample size range between 1,293 to 5,016. All measures are normally or modest normally distributed as indicated in **Supplementary table 1**.

#### Supplementary Note 2: Description of mediation models.

The SEM for this mediation model for the *i* th subject ($1 \leq i \leq n$) is given by:

$$z_i = \beta_0 + \beta_{xz} x_i + \varepsilon_{zi},$$
$$y_i = \gamma_0 + \gamma_{zy} z_i + \gamma_{xy} x_i + \varepsilon_{yi}$$

It is posited that the error terms ($\epsilon z_i$, $\epsilon y_i$) are uncorrelated, a critical presumption for causal inference when conducting mediation analysis. The assumption of multivariate normality for the error terms is also made, as it is an essential precondition for defining direct, indirect, and total effects. It should be highlighted that the two structural equations are interconnected, and the inference drawn from them is concurrent, rather than from two separate standard regression equations. More information can be found here [1].

#### Mediation analyses

We conducted mediation analyses (Baron & Kenny, 1986; Preacher & Kelley, 2011) using the lavaan package for R to examine the direct and indirect effects of the prediction from genetic predisposition (quantified as the PGSs of educational attainment, cognitive and noncognitive skills) and manifestation of variation in academic achievement (Figure 1).

The mediation model estimates the indirect effect of the predictor (X) on the outcome (Y) via a mediator, i.e., an intervening variable (mediator; M; in this project, the single-timepoint or developmental environmental composite) by regressing M on X and regressing Y on both X and M using two separate equations:

$$1)\ M_i = d_{M.X} + aX_i + e_{M.Xi}$$

Where Mi is the mediator for individual I; dM.X is the intercept for the mediator (M); aXi is the slope of M regressed on the predictor (X) and eM.Xi is the measurement error for individual i.

$$2)\ Y_i = d_{Y.MX} + bM_i + c'X_i + e_{Y.MXi}$$

Where Yi is the outcome for individual I; dY.MX is the intercept for the outcome (Y); bMi is the slope of the outcome (Y) regressed on the mediator (M) controlling for the predictor (X); c'Xi is the slope of the outcome (Y) regressed on the predictor (X) controlling for the mediator (M) and eY.MX is the measurement error for individual i.

The indirect effect of the predictor on the outcome (i.e., the mediation effect) is defined by a^b^, with the sample estimate signified by the circumflex ("^").

When $a^{\wedge} \times b^{\wedge} = c^{\wedge} - c^{\wedge\prime}$ , then $c^{\wedge} = a^{\wedge} \times b^{\wedge} + c^{\wedge\prime}$. Implementing SEM allows for, a^ and b^ can be derived simultaneously and for testing more complex models with latent class predictor, outcomes, and mediators.

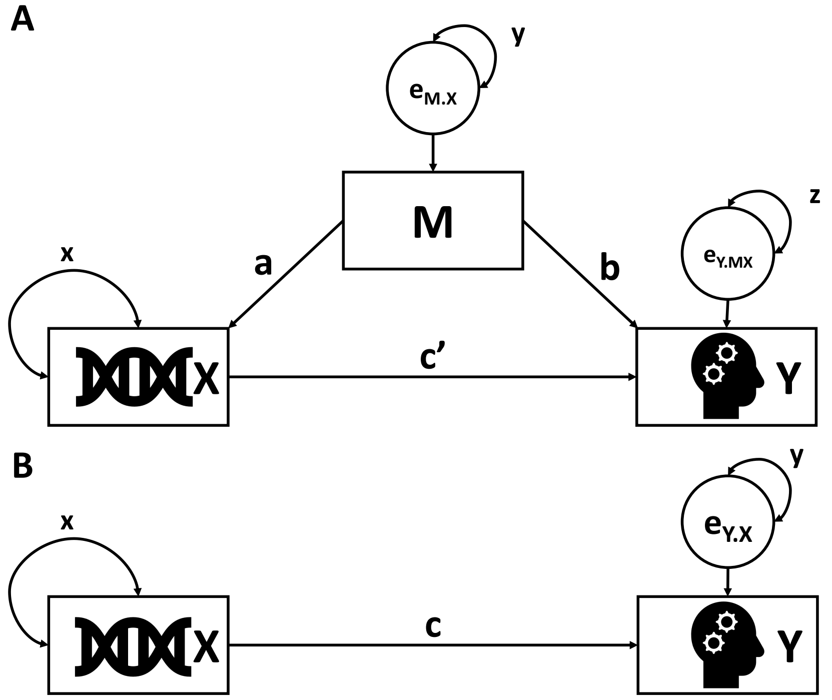


Note: Mediation model of the DNA (X) of cognitive/educational outcomes (Y) that is (panel A) versus is not (panel B) mediated by M, which denotes the noncognitive skills, either single-timepoint or developmental using time-lagged data. Circles indicate residuals. Parameters a, b and c represent regression weights. Parameters x, y and z represent variance parameters.

### Supplementary Figures

#### Supplementary Figure 1: Correlation matrix between PGS and academic achievement.

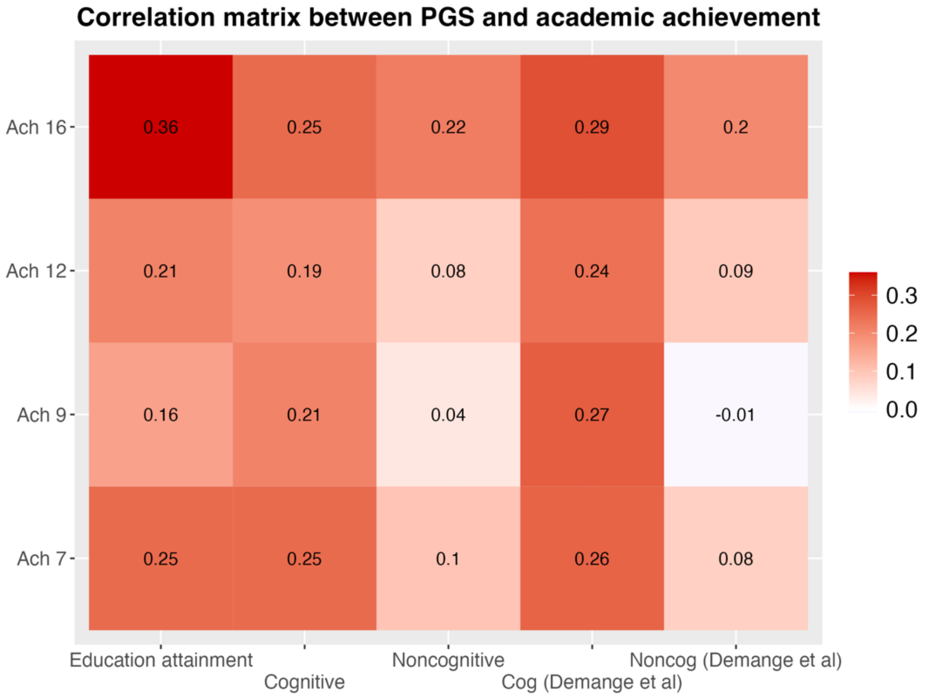


Note: Ach 7: A composite mean score of academic performance at age 7; Ach 9: A composite mean score of academic performance at age 9; Ach 12: A composite mean score of academic performance at age 12; Ach 16: Mean of the grade obtained across the GCSE subjects (English, Maths, and Science); EA: Educational attainment: Educational attainment polygenic score; Cognitive: Cognitive polygenic score; Noncognitive: Non-cognitive polygenic score; Cog (Demange et al): Cognitive polygenic score derived from previous work by Demange et al., 2021; Noncog (Demange et al): Non-cognitive polygenic score derived from previous work by Demange et al., 2021.

**Supplementary Figure 2: Correlations between latent factors of non-cognitive skills, academic achievement and education and cognition-associated polygenic scores.**

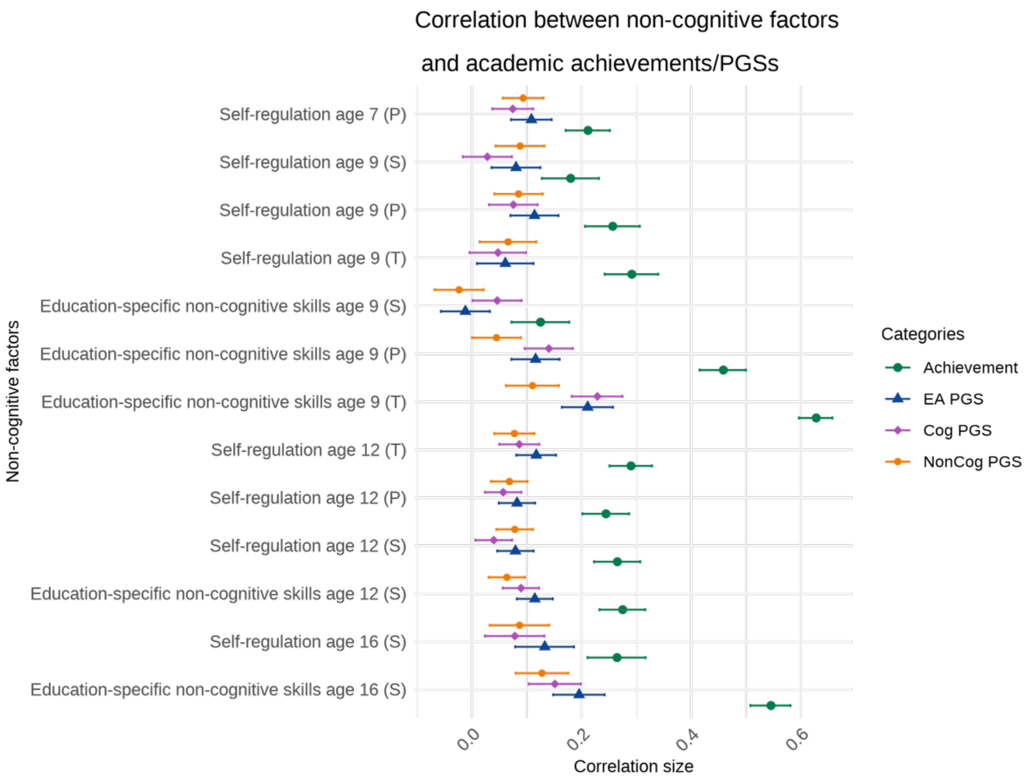


Note: Data on non-cognitive skills was collected from different raters: (P) = parent-reported, (T) = teacher-reported and (S) = Self-reported. Academic achievement was measured contemporaneously with each non-cognitive factor. EA PGS[2] = educational attainment polygenic score. Cog PGS[3] = cognitive skills polygenic score. NonCog PGS[3] = non-cognitive skills polygenic score. Each dot indicates the size of the correlation coefficient, and error bars indicate 95% confidence intervals. Sample size for each correlation analysis is provided in Supplementary Table 2.

**Supplementary Figure 3: Comparison of indirect effects of educational attainment prediction between g-uncorrected/ g-corrected noncognitive skills using two-mediators model.**

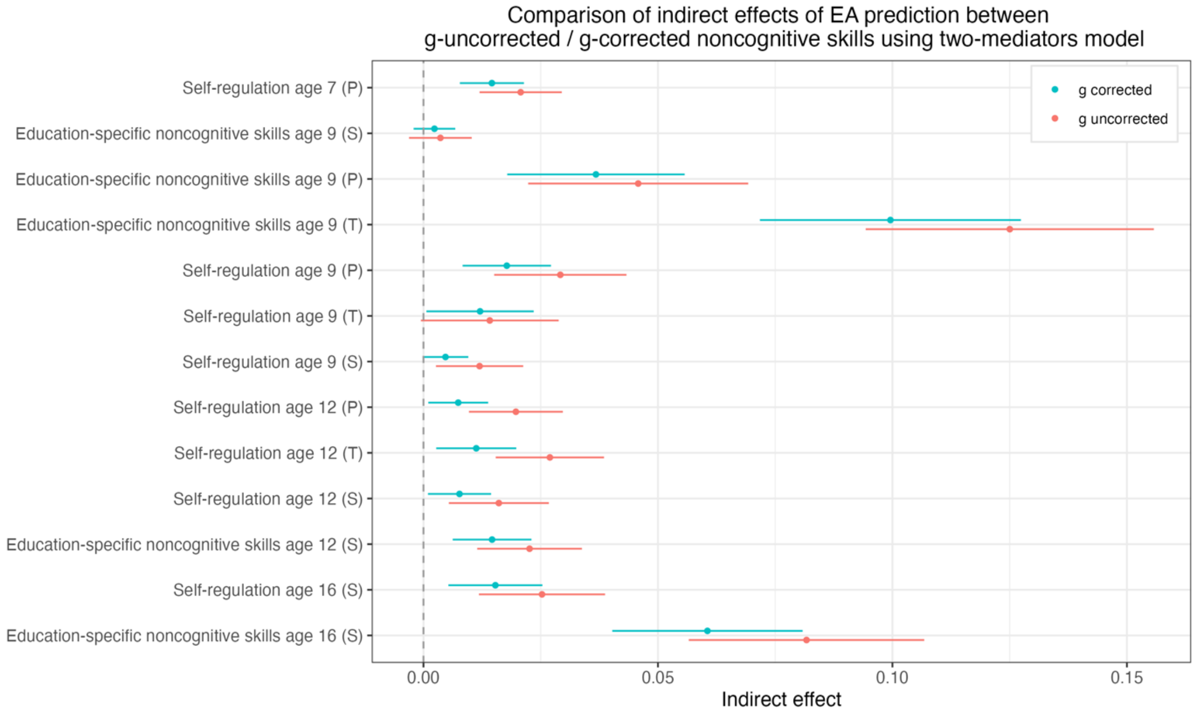

**Supplementary Figure 4: Comparison of indirect effects of cognitive prediction between g-uncorrected/ g-corrected noncognitive skills using two-mediators model.**

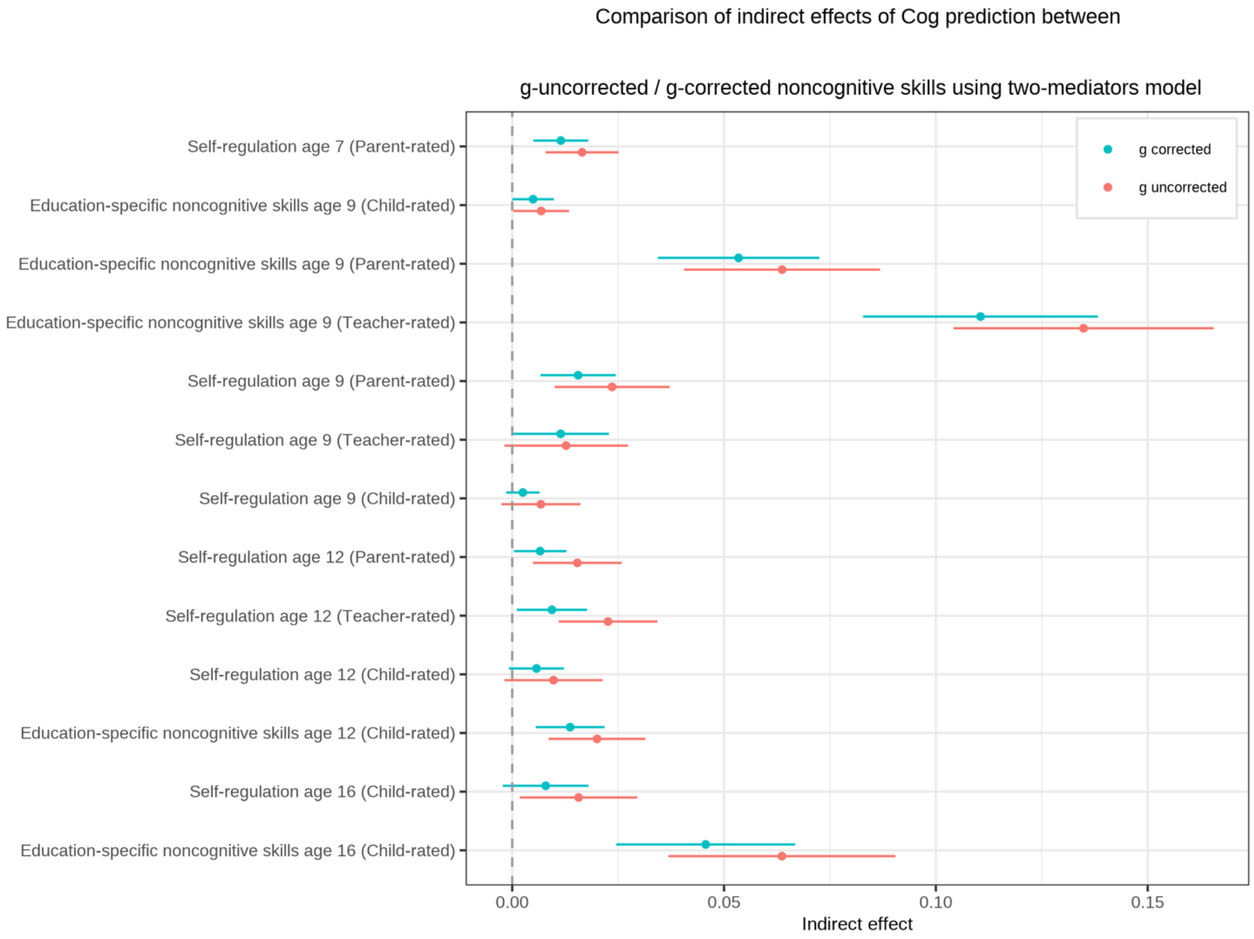

**Supplementary Figure 5: Comparison of indirect effects of noncognitive prediction between g-uncorrected/ g-corrected noncognitive skills using two-mediators model.**

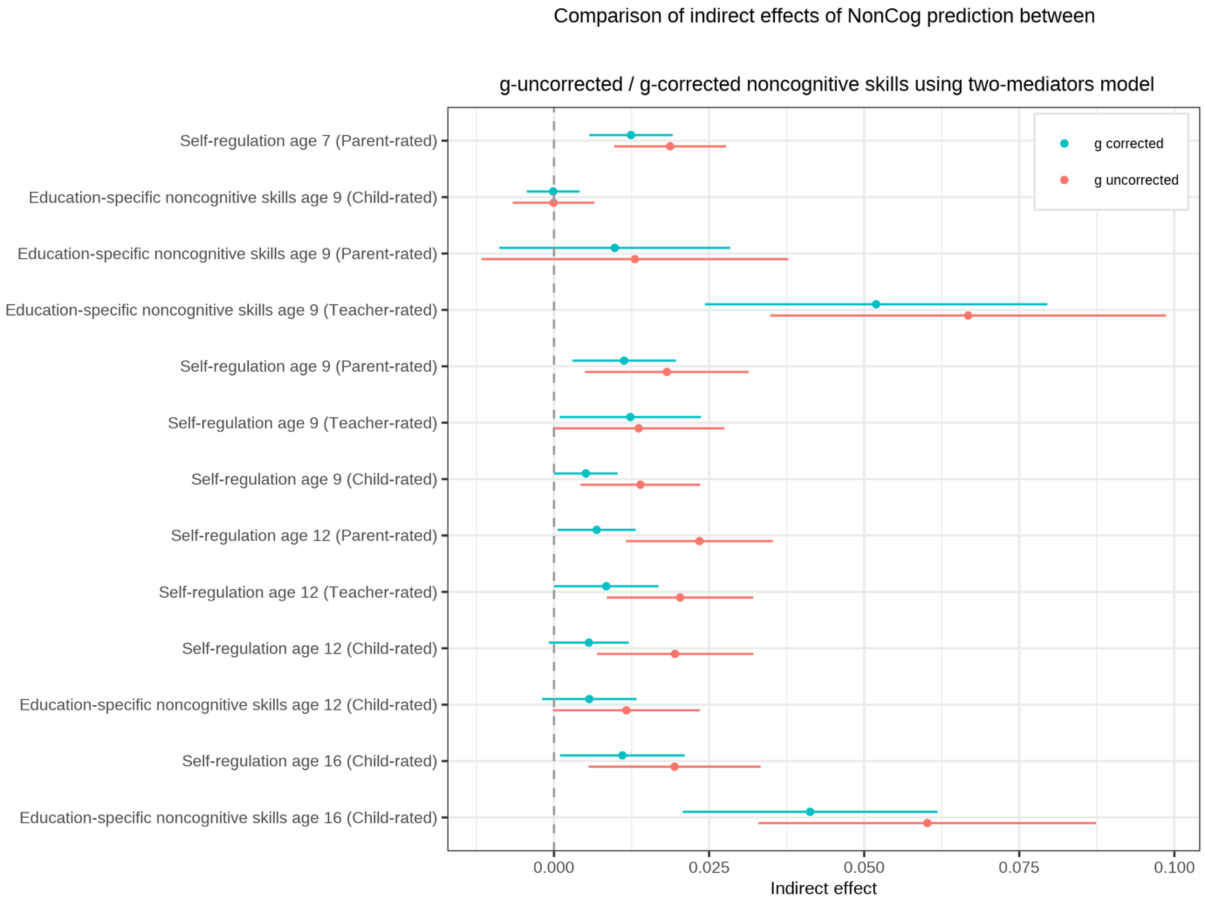

**Supplementary Figure 6: Comparison of indirect effects of educational attainment prediction between SES-uncorrected/ SES-corrected noncognitive skills using two-mediators model.**

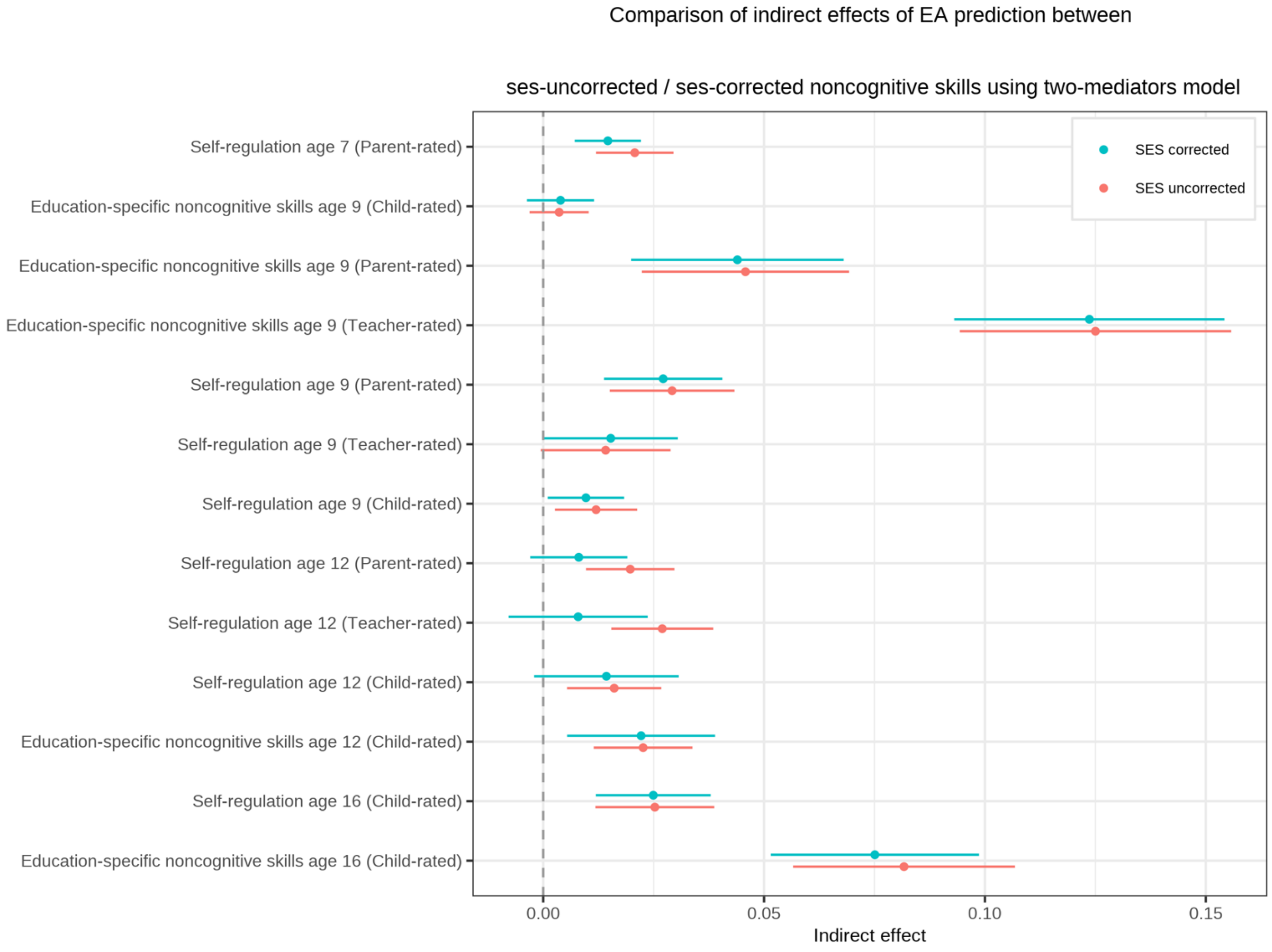

**Supplementary Figure 7: Comparison of indirect effects of cognitive prediction between SES-uncorrected/ SES-corrected noncognitive skills using two-mediators model.**

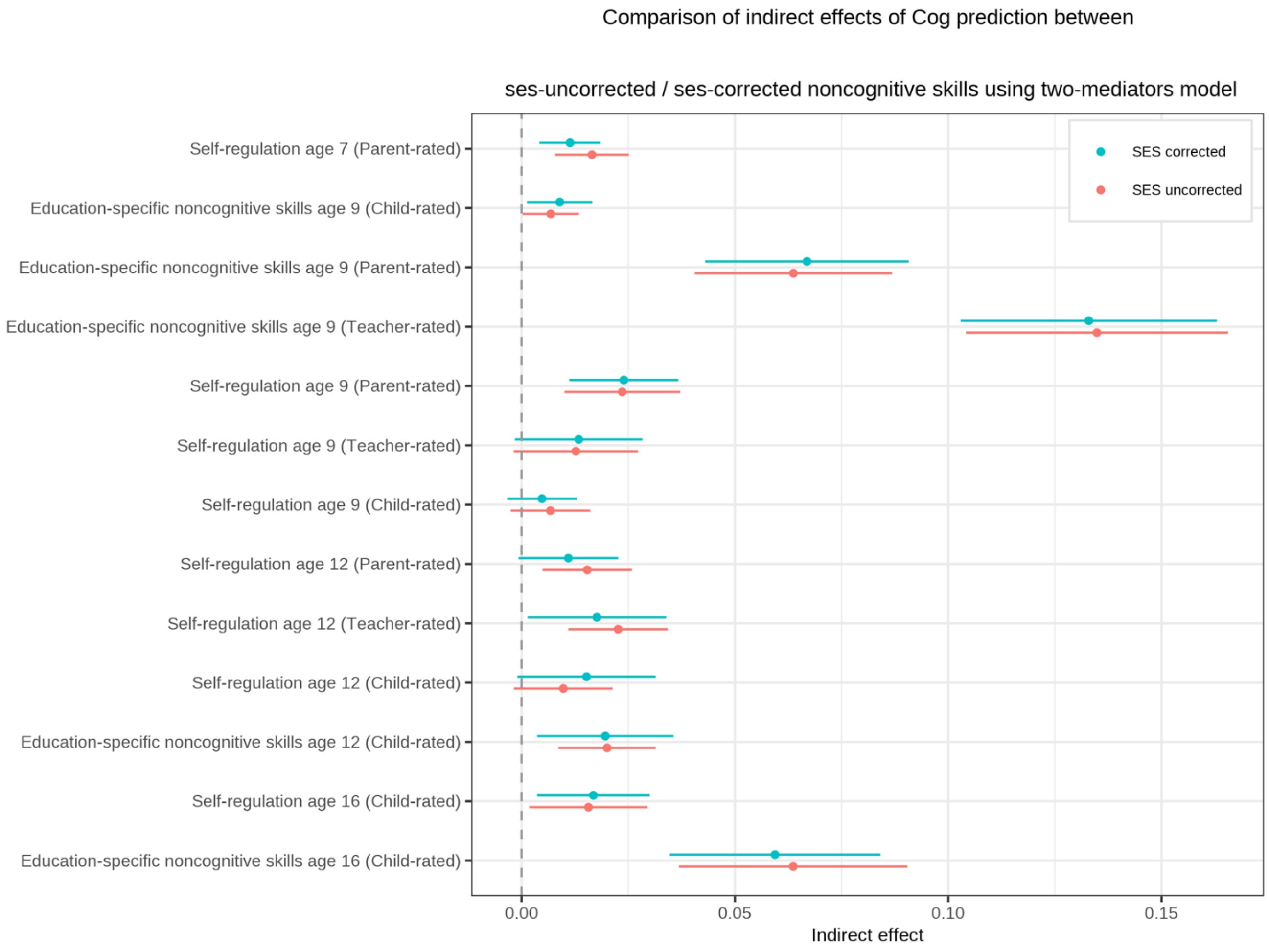

**Supplementary Figure 8: Comparison of indirect effects of noncognitive prediction between SES-uncorrected/ SES-corrected noncognitive skills using two-mediators model.**

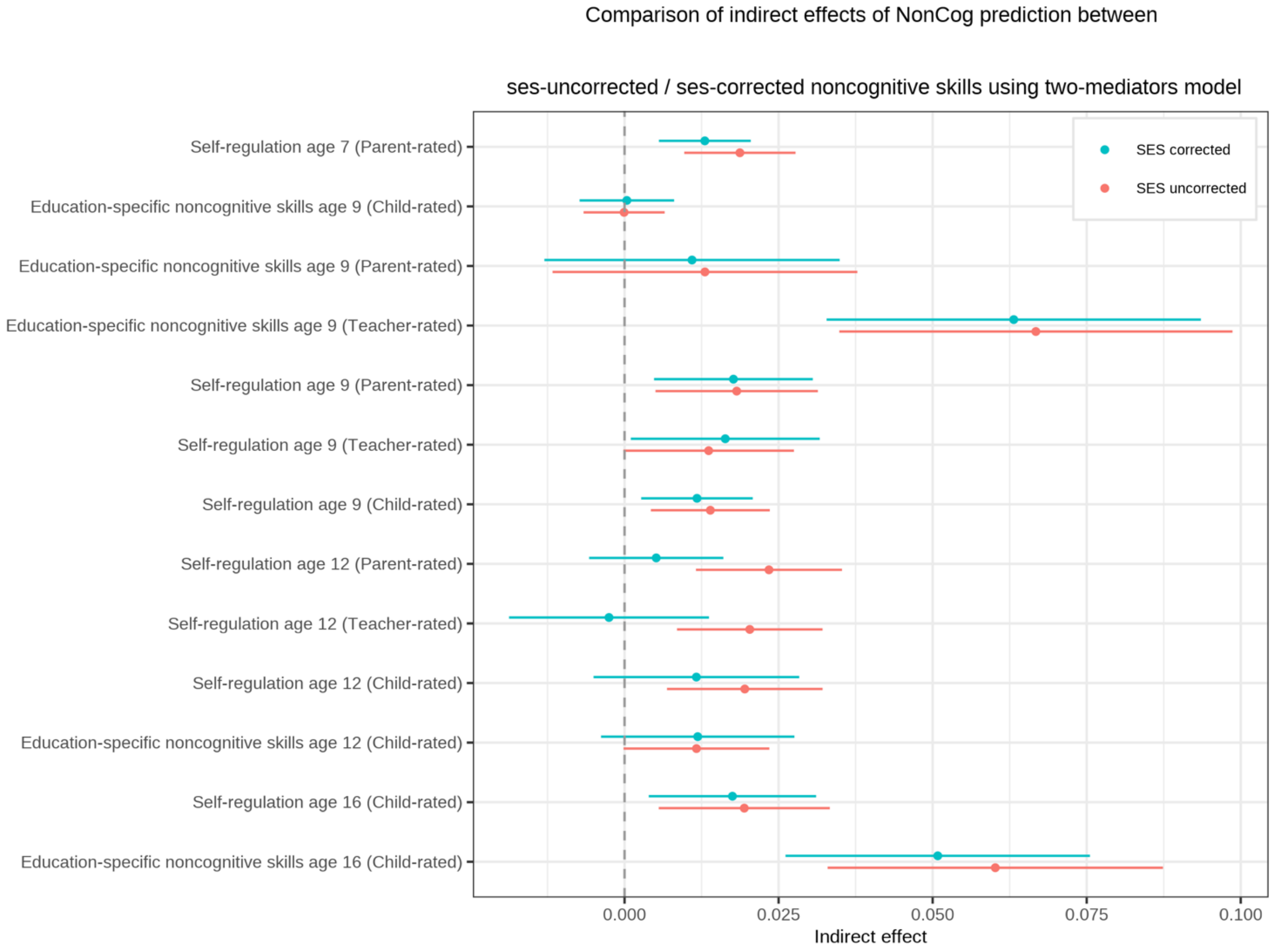
